# Deviant functional connectivity patterns in the EEG related to developmental dyslexia and their potential use for screening

**DOI:** 10.1101/2025.03.04.641551

**Authors:** Yaqi Yang, Zhaoyu Liu, Brian Wong, Shuting Huo, Jie Wang, Tan Lee, Fumiko Hoeft, Urs Maurer

## Abstract

Developmental dyslexia (DD) is a common learning disorder with potential neural origins. While EEG-based brain activation measures combined with machine learning have shown promise for DD screening, these approaches often lack validation on independent participants—a crucial step for practical application. This study developed an EEG-based screening approach and investigated the neural correlates of DD in Chinese children. EEG signals were recorded from 130 children (82 with DD, 48 typically developing) aged 7–11 during resting-state and working memory tasks. The EEG data were preprocessed into clean segments to compute functional connectivity (FC) matrices across four frequency bands (delta, theta, alpha, beta). The segments were split into two independent samples, ensuring independence at the participant level: Sample 1, used for training and five-fold cross-validation of the convolutional neural networks, and Sample 2, used for cross-sample validation with the trained model. The beta-band FC index in the eyes-open condition achieved the highest within-sample classification accuracy (98%) and cross-sample accuracy (70%, p < .001). Discriminative FC patterns revealed that children with DD exhibited reduced temporal-parietal and central connectivity but increased frontal-central connectivity, likely reflecting compensatory mechanisms. Within the DD group, stronger FCs showed significant negative correlations with Chinese word reading accuracy and fluency. These findings suggest that EEG-based FC measures can effectively distinguish DD and reveal neural markers associated with impaired reading performance. This approach shows promise for non-invasive screening and deeper insight into the neural basis of DD, particularly in non-alphabetic language systems.

## 1. Introduction

Developmental dyslexia (DD) is a specific learning disability characterized by persistent difficulties in word decoding despite adequate instruction, intelligence, and unimpaired sensory abilities (Démonet et al., 2004; Lyon, Shaywitz, & Shaywitz, 2003). Representing one of the most prevalent learning disabilities, DD affects approximately 5-10% of children globally (Shaywitz et al., 2008). Even though DD is thought to be caused by an underlying deficit in phonological processing (Ramus, 2003), a similar prevalence rate has been reported for children in Hong Kong learning to read Chinese, where the role of phonological processing in reading is distinct (Chan et al., 2007). Children with DD may face not only a higher risk of difficulties in academic performance but also an elevated risk for a range of emotional difficulties, such as anxiety (Xiao et al., 2022), depression (Alexander-Passe, 2012), and low self-esteem (Novita, 2016). Long-term studies have suggested that these DD-related difficulties persist into adulthood (Roitsch & Watson, 2019). To counter these adverse outcomes, accurately and promptly screening children with DD and understanding the neural substrates of the disorder are essential.

Accurate screening of children with DD is crucial to effective research and intervention. The diagnostic criteria for DD are primarily based on the standards outlined in the Diagnostic and Statistical Manual of Mental Disorders, 5th Edition (DSM-5) (American Psychiatric Association, 2013). Traditional diagnostic procedures for DD mainly involve a battery of behavioral assessments designed to evaluate a child’s cognitive, linguistic, and academic skills. These assessments typically include the core measures of word decoding accuracy and fluency, additional literacy-related measures such as comprehension and phonological awareness, and measures of basic intellectual performance (Schulte-Körne, 2010; Chung & Ho, 2010). Children scoring among the lowest 10% of the norm population in word reading tests (Lyytinen et al., 2015; Price et al., 2022), with average intelligence, and without other perceptual deficits or mental and neurological disorders are diagnosed as having DD (Di Folco et al., 2022; O’Brien & Yeatman, 2021).

However, the reliability of DD diagnosis based on behavioral assessments is limited by the natural variation in reading progress within the first 1-2 years among typically developing (TD) children. This variability can lead to underdiagnosis for at-risk children during the critical early phase of literacy acquisition. For example, one study showed that 13.4% of children at risk in kindergarten were not diagnosed as reading disabled until grade 5 (Catts et al., 2012). Given DD’s assumed neurological origin, neuroimaging studies have tried to identify differential neural patterns between children with DD and TD children. These studies reported consistent neural abnormalities primarily within the left hemisphere’s language network, particularly in the temporoparietal and occipitotemporal regions and the inferior frontal gyrus (Beelen et al., 2019; Yan et al., 2021). Based on these neural abnormalities, studies integrating neuroimaging data with machine learning have shown promise in DD screening, achieving remarkable classification accuracy in distinguishing individuals with dyslexia (Cui et al., 2016; Tamboer et al., 2016; Tanaka et al., 2011). Additionally, these abnormalities can be used to predict the long-term reading ability of children with DD (Hoeft et al., 2011). Therefore, neuroimaging-based screening methods are instrumental in identifying children with DD and discovering associated neural biomarkers. These biomarkers could potentially be used to predict future reading outcomes and assist in the early detection of children at risk for DD, offering a comprehensive approach to tackling this complex disorder.

Electroencephalogram (EEG), a readily accessible, cost-effective, and non-invasive neuroimaging technique, could be crucial in discerning neural patterns and discriminating brain patterns for DD screening. Parallel to fMRI studies, EEG studies have illuminated the brain abnormalities associated with DD at rest and during cognitive tasks. Researchers have utilized frequency-domain features, such as spectral power density and spectral centroid, to examine the electrophysiological abnormalities in children with DD (Papagiannopoulou & Lagopoulos, 2016; Ribeiro et al., 2020). For example, abnormalities in beta-band activity (12 to 30 Hz) have been observed in the frontal and central brain areas of children with DD (Eroğlu et al., 2022; Spironelli et al., 2008), which are linked to executive functions and visuomotor integration, suggesting a broader impact on cognitive processing in DD (Basharpoor et al., 2022).

In addition to beta frequency band abnormalities, previous studies have attempted to identify electrophysiological abnormalities in children with DD by examining a wide range of EEG features. These include a broad spectrum of features, such as time-domain features like event-related potential (ERP) amplitude and temporal skewness (Eroğlu et al., 2022; Hämäläinen et al., 2013), complexity features such as approximate entropy and fractal dimension (Pappalettera et al., 2022; Lau et al., 2022), connectivity features like Pearson correlation coefficients (PCC) and phase locking value (PLV) (Bosch-Bayard et al., 2020; Granados et al., 2021), and graph metric features like degree and betweenness centrality (González et al., 2016; Xue et al., 2020). Studies have been directed at integrating these EEG features with machine learning algorithms to facilitate screening children with

DD. The majority of these studies have predominantly employed conventional machine learning techniques such as Support Vector Machine (SVM), Logistic Regression (LR), and Random Forest (RF). These efforts have yielded classification accuracy rates ranging from 70% to 96% (Frid & Manevitz, 2020; Gallego-Molina et al., 2022; Ortiz et al., 2020; Parmar & Paunwala, 2022; Rezvani et al., 2019). While some of these studies have reported impressive accuracy rates, they often relied on small sample sizes and were predominantly focused on alphabetic languages.

To the best of our knowledge, there is a lack of EEG-based DD classification studies that employ cross-validation on new, independent samples. This is a critical point, as pointed out by Poldrack et al. (2020). Machine learning methods often utilize experimental protocols that involve subdividing samples into distinct training and test datasets for learning and cross-validation. These protocols also include repeatedly assigning data to training and test sets through multiple cycles or folds. However, this approach can bear the problem of test data leakage and non-independence between training and test sets, potentially leading to model overfitting and poor generalization. Poldrack et al. (2020)emphasized the necessity of performing cross-validation on a separate, isolated dataset rather than within the original training sample to obtain a genuine measure of predictive accuracy. Generalizing a new sample is a necessary precondition if a predictive algorithm is applied in practice to identify new children with DD. Therefore, comprehensive research that includes a broader spectrum of language systems and encompasses larger sample sizes is needed. Furthermore, such research should rigorously apply cross-sample validation techniques in EEG-based deep learning frameworks for screening DD in children.

Unlike traditional machine learning methods, which often require explicit feature extraction, Convolutional Neural Networks (CNNs) autonomously learn to extract complex features through their layered architecture when trained on substantial datasets (Chauhan et al., 2018; Wu et al., 2017). CNNs is specially designed to learn hierarchical representations of input data through convolutional and pooling layers. The convolutional layers use adjustable filters on input data and, in conjunction with pooling layers that reduce the spatial dimensions while preserving essential information, enable the network to progressively learn more complex features through its layered structure (Albawi et al., 2017; Sun et al., 2017). Successful applications of CNNs using structured EEG data have been seen in the screening of neural developmental disorders such as Autism Spectrum Disorder (ASD) (Ari et al., 2022; Peya et al., 2020), Attention Deficit Hyperactivity Disorder (ADHD) (Chen, Song, & Li, 2019; Moghaddari, Lighvan & Danishvar, 2020), and Alzheimer’s disease (Lopes et al., 2023; Wen et al., 2020).

Incorporating EEG functional connectivity (FC) measurements can enhance the efficiency of CNNs in screening for neurodevelopmental disorders (Fu et al., 2016). FC quantifies the synchronization between pairs of EEG electrodes over time or phase, allowing for the creation of a matrix that resembles an image for each electrode pair (Fraschini et al., 2016). This transformation fully capitalizes on the strengths of CNNs, whose architecture is inspired by the visual cortex and is particularly adept at processing graph-like structures. Consequently, a hybrid approach that combines EEG-based FC measurements with CNNs could significantly improve the screening process for children with DD. This combined tool has the potential to provide a deeper understanding of the neurophysiological underpinnings of DD, paving the way for more accurate and efficient EEG-based screening techniques.

While previous studies have shown promise in classifying children with DD, several limitations persist within these studies. A primary concern is that most EEG-based DD screening studies have relied on relatively small sample sizes, typically ranging from 11 to 29 individuals with DD (Formoso et al., 2021; Mahmoodin et al., 2016; Ortiz et al., 2020; Zainuddin et al., 2016; Zainuddin et al., 2018). Furthermore, there is a lack of cross-sample validation using independent samples, which is necessary to address the high variability observed between and within subjects. This lack of extensive validation undermines the generalizability of the classification outcome (Poldrack et al., 2020). Additionally, only a few of these EEG-based DD classification studies have attempted to uncover potential neural biomarkers contributing to the classification of DD. To our knowledge, no previous study has explored the behavioral association between the identified biomarkers and reading ability. This dimension could shed new light on DD’s underlying mechanisms and potential interventions. Moreover, the existing research has primarily focused on DD classification within alphabetic language contexts. DD’s cognitive and neural manifestations can differ significantly when considering non-alphabetic languages like Chinese (Wong et al., 2012; Su et al., 2015). Therefore, developing and adapting models specifically tailored to DD’s unique profiles in non-alphabetic languages is imperative. Lastly, previous studies have not systematically explored the differences between resting-state and task-state EEG measures in screening children with DD. Investigating this could help determine whether DD is a domain-general or domain-specific neurodevelopmental disorder.

To overcome the limitations of existing research, we proposed a novel approach in the current study that integrated EEG-based FC measures both at resting- and task-states with CNNs for effective DD screening and identification of behaviorally relevant neural biomarkers. Our process began with the acquisition and pre-processing of raw EEG data. We then transformed the pre-processed EEG signals into FC matrices using three distinct measures: Pearson Correlation Coefficients (PCC), Phase Locking Value (PLV), and Rho Index (RHO). These measures were computed across four frequency bands—delta, theta, alpha, and beta—under four experimental conditions: eyes-open and eyes-closed resting states, as well as one-back and two-back working memory tasks with Chinese characters. The resulting matrices were randomly divided into two independent samples for each FC measurement, frequency band, and experimental condition. The first sample served as the training and validation set for the CNNs model, while the second was designated for cross-validation to affirm the model’s robustness and generalizability across the participant level. This approach allowed us to evaluate each FC measure thoroughly and examine the effectiveness of each frequency band and task condition in identifying children with DD. Furthermore, we identified discriminative FCs that significantly contributed to the classification of DD. Lastly, we examined the correlations between these discriminative FCs and Chinese word reading abilities within DDs and TDs separately. Our findings may enhance the understanding of Chinese DD’s neurophysiological bases and refine screening approaches.

## 2. Methods

### 2.1 Participants

A total of 130 participants, including 82 children with DD and 48 TD children in the second or third grade (aged 7 to 11 years, shortly after the children with DD received a formal diagnosis), were recruited in our current study. All children in the study were part of a larger research project approved by The Joint Chinese University of Hong Kong—New Territories East Cluster Clinical Research Ethics Committee (The Joint CUHK-NTEC CREC). The children were native Cantonese Chinese speakers recruited from primary schools and education authorities in Hong Kong, and written consent was obtained from both, the children and their guardians.

To be included in the study, children with DD had to meet the following criteria: (1) a formal diagnosis of DD by an educational or clinical psychologist based on the Hong Kong Test of Specific Learning Difficulties in Reading and Writing for Primary School Students—Third Edition [HKT-P(III)] (Ho et al., 2007), which required adequate IQ (higher than 85) and poor literacy (−1 SD or below), and at least one area of cognitive-linguistic deficit (−1 SD or below) (Chung, 2017); and (2) no history of brain injury, birth complication, significant sensory impairment, or other neurological or psychological disorders (e.g., ADHD). TD children were identified based on their parents reporting that the child had no reading or writing difficulties. Table 1shows the demographic information of the two groups, who did not differ significantly in age, maternal or paternal education level, or family income but differed significantly in Raven’s non-verbal intelligence, Chinese word reading accuracy, and character and word reading fluency.

**Table 1.** Demographic table of the typically developing (TD) and developmental dyslexia (DD) groups.

| Measurement | TD | DD | t |
| --- | --- | --- | --- |
| Male-to-female ratio | 24 : 24 | 41 : 41 | - |
| Age [years] | 8.02 ($\pm 0.53$) | 7.99 ($\pm 1.35$) | -0.16 |
| Raven's non-verbal intelligence [raw scores] | 27.96 ($\pm 3.14$) | 24.70 ($\pm 6.65$) | 3.13** |
| Family income [category]<sup>1</sup> | 4.09 ($\pm 1.44$) | 3.90 ($\pm 1.52$) | 0.67 |
| Maternal education [level]<sup>2</sup> | 3.22 ($\pm 1.52$) | 3.34 ($\pm 1.48$) | -0.45 |
| Paternal education [level]<sup>2</sup> | 3.24 ($\pm 1.54$) | 3.15 ($\pm 1.70$) | 0.30 |
| Chinese word reading accuracy [correct items] | 89.40 ($\pm 17.57$) | 48.43 ($\pm 25.23$) | 9.82** |
| Chinese character and word reading fluency [number of correct items per minute] | 151.28 ($\pm 49.00$) | 61.37 ($\pm 36.50$) | 11.74** |

**Notes:** <sup>1</sup>Monthly family income was categorized as follows: 1 for HKD 10,000 (USD 1,280) or below, 2 for HKD 10,001–20,000 (USD 1,281–2,560), 3 for HKD 20,001–30,000 (USD 2,561–3,840), 4 for HKD 30,001–40,000 (USD 3,841–5,120), 5 for HKD 40,001–50,000 (USD 5,121–6,400), and 6 for HKD 50,001 (USD 6,401) or above.
<sup>2</sup>Maternal and paternal educational levels were coded with the following scale: 1 for middle school or below, 2 for high school, 3 for preparatory school, 4 for college, and 5 for postgraduate studies.
\*\*Significant at the .01 level (2-tailed).

### 2.2 Behavioral assessments

Chinese word reading accuracy (Wang et al., 2020): The evaluation of Chinese word reading accuracy used a list of two-character Chinese words derived from the Hong Kong Corpus of Primary School Chinese (Leung & Lee, 2002). The second character of each word progressively increased in difficulty. Children were asked to read the second word aloud and were awarded a point for each correct response. The task was concluded after the participant provided ten consecutive incorrect responses. The total number of correct responses was regarded as the score for Chinese word reading accuracy.

Chinese character and word reading fluency (Siu et al., 2018): Reading fluency was measured using three sub-tests. The first two sub-tests involved presenting lists of Chinese single-character words, classified as either consistent or inconsistent with the phonetic radical. The third sub-test presented a list of two-character words. Children were tasked with reading the items as quickly and accurately as possible. The reading fluency score for each sub-test was quantified by the count of items correctly read by the participants within a one-minute interval. The sum of the three sub-test scores was the Chinese character and word reading fluency score.

### 2.3 EEG assessments

The eyes-open and eyes-closed resting-state paradigm: During the EEG recording session, children were first directed to participate in two distinct three-minute resting-state paradigms, with their EEG activity being recorded throughout: an eyes-open condition and an eyes-closed condition. Initially, children were guided to sit comfortably and relax. In the eyes-open condition, they were asked to fixate on a cross displayed on a computer screen for three minutes without being engaged in any specific tasks. Conversely, for the eyes-closed condition, children were instructed to look straight ahead, close their eyes, and remain motionless for three minutes.

The verbal *n*-back tasks (Wang et al., 2022): Children’s verbal working memory was assessed using *n*-back tasks with concurrent EEG recording. Chinese character stimuli were presented sequentially, and the children either needed to detect immediate (one-back) repetitions or characters presented two trials earlier (two-back). Working memory load is higher in the two-back task than in the one-back task (Haatveit et al., 2010). Sixty non-target and 20 target stimuli were presented in each of the two tasks. The characters were arranged to avoid apparent semantic relatedness or orthographic similarity between two consecutive stimuli except for the targets in the one-back task. Each character was displayed for 500 ms, followed by a 3500-ms fixation (see Figure 1 for the trial procedure). Children were instructed to press the “1” key as accurately and quickly as possible when they detected a target, and no response was needed for non-targets. Before the formal experiment, children completed a practice block with 14 characters to ensure understanding. The practice block could be repeated if needed.

**Figure 1.**
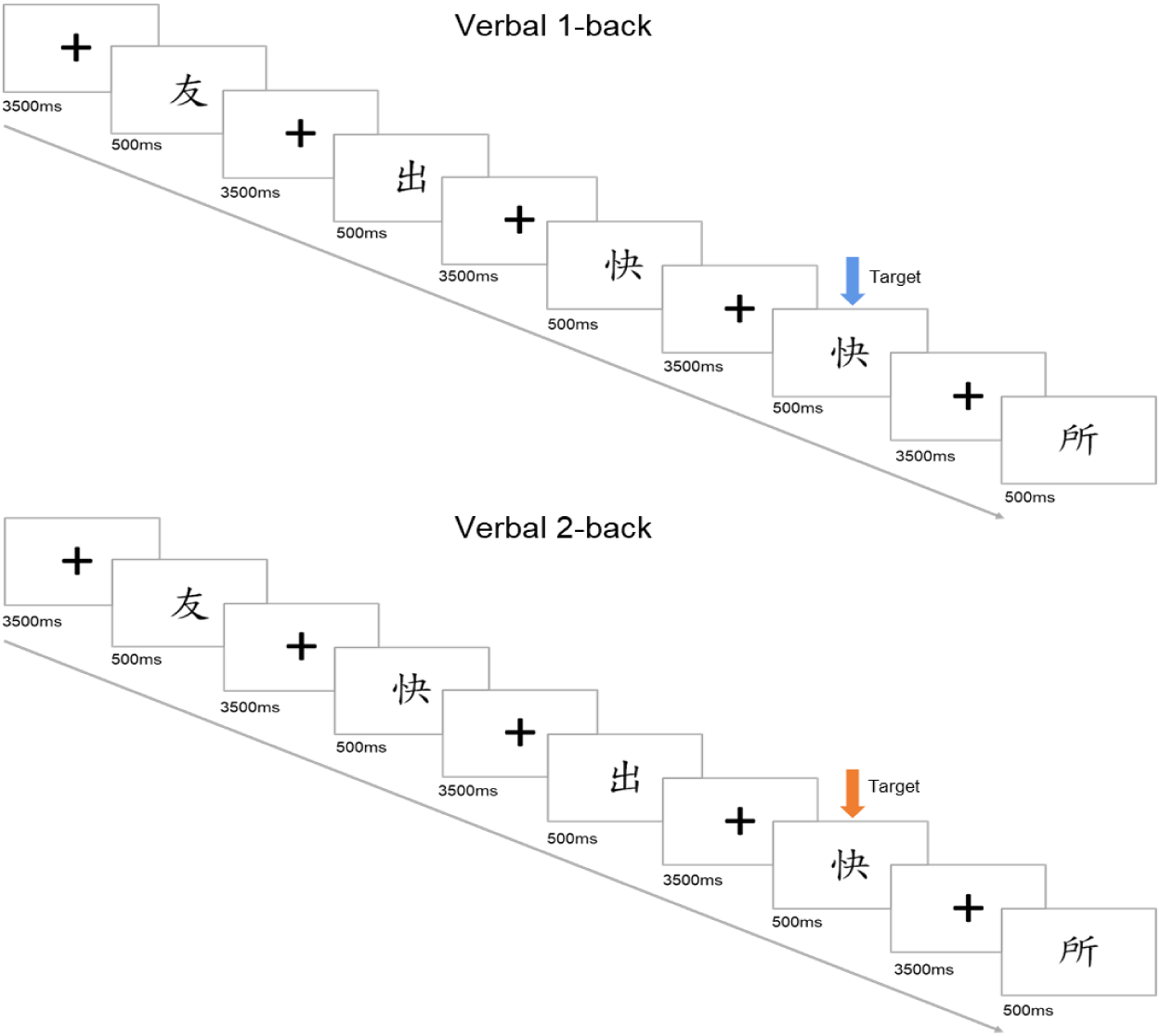
**Example trials in the *n*-back tasks assessing verbal working memory.**

### 2.4 The flowchart of this study

The current study proposed an EEG-based deep-learning classification framework for screening DD. Initially, we collected and pre-processed raw EEG signals into cleaned 2-second segments. These segments were subsequently transformed into delta, theta, alpha, and beta frequency bands via the Fast Fourier Transform (FFT) (Brigham, 1988). Following this, we computed FC matrices for each segment within each frequency band in the four experimental conditions (i.e., eyes-open, eyes-closed, verbal one-back, and verbal two-back conditions). The segments were randomly divided into two independent samples: Sample 1 and Sample 2. Sample 1 was utilized to train and validate the CNNs model, while Sample 2 was reserved for cross-sample validation. The trained CNNs model, using the optimal frequency band FC indices, was employed to predict DD in Sample 2. A permutation test was conducted to determine whether the cross-sample validation accuracy exceeded the chance level. Further, independent-sample *t*-tests with False Discovery Rate (FDR) correction (Genovese et al., 2002) were performed to identify discriminative FCs contributing to the DD classification in Sample 1. Finally, separate correlation analyses were conducted to explore the associations between these discriminative FCs and Chinese word reading abilities in children within the DD and TD groups, respectively. Figure 2 shows the flowchart of the current study.

**Figure 2.**
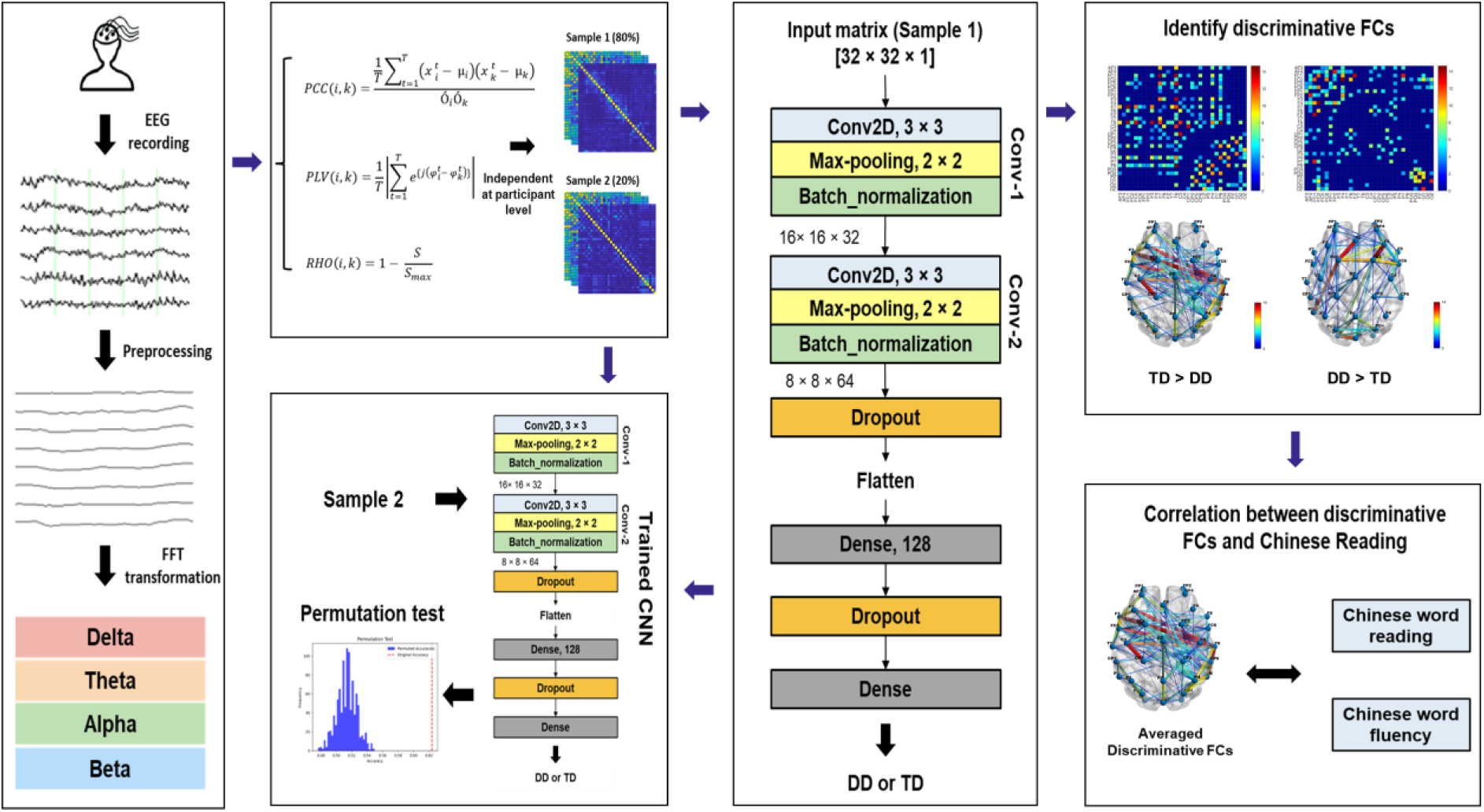
The flowchart of the current study. Notes: Abbreviations: EEG (Electroencephalography), FFT (Fast Fourier Transform), FC (Functional Connectivity), PCC (Pearson Correlation Coefficient), PLV (Phase Locking Value), RHO (Rho Index), CNN (Convolutional Neural Networks), DD (Developmental Dyslexia), TD (Typically Developing).

### 2.5 EEG recording and pre-processing

Both resting-state and task-state EEG data were recorded using the 128-channel EGI system (Electrical Geodesics, Inc.) at a sampling rate of 500 Hz, with recordings referenced to Cz and an online filter of 0.1-100 Hz. Impedances were maintained below 50 kΩ. The data were then pre-processed using the BrainVision Analyzer (https://www.brainproducts.com/solutions/analyzer/)filtered offline with a 0.3-70 Hz bandpass filter and a 50 Hz notch filter. Channels identified as bad were excluded, and the remaining EEG signals underwent independent component analysis (ICA) to correct eye movement artifacts. These excluded channels were subsequently spline interpolated and all channels re-referenced to the average reference. Afterward, we segmented the continuous resting-state EEG recordings into 2-second segments as input for the CNNs. For the verbal *n*-back tasks, we extracted only the last 2 seconds of the fixation period following the non-target as the 2-second segments.

Portions of the EEG data from the working memory tasks and eyes-closed resting state have previously been analyzed to explore group differences in EEG band power between children with DD and TD children, as reported in the study by Wang et al. (2022). The pre-processing protocol in the current study parallels that of Wang et al. (2022), except for artifact rejection procedures. To streamline computations and mitigate volume conduction effects from adjacent channels (Kashefpoor et al., 2016), we selected those 32 channels that corresponded to the 10-20 system from the original 128-channel EGI system following the standard transformation criteria (Luu & Ferree, 2005). These channels included AF3, AF4, FC1, FC2, FC5, FC6, FP1, FP2, F3, F4, F7, F8, FZ, T7, T8, C3, C4, CP1, CP2, CP5, CP6, CZ, P3, P4, P7, P8, PO3, PO4, PZ, O1, O2, and OZ. Artifact rejection was performed on the selected channels; segments exhibiting amplitudes beyond ±100 μV were discarded.

The retained segments were then prepared for further analysis. Fast Fourier Transform (FFT) (Brigham, E. O., 1988) was applied to convert the EEG data from the time domain to the frequency domain, followed by bandpass filtering to obtain delta (0.5-3.5 Hz), theta (4-8 Hz), alpha (8-12 Hz), and beta (12-30 Hz) frequency bands. In total, we preserved 5,594 segments for the eyes-open condition and 6,204 segments for the eyes-closed condition, including 2,010 and 2,330 segments from TD children, respectively. In the task states, there were 4,245 segments from the verbal one-back condition and 4,114 segments from the verbal two-back condition, including 1,765 and 1,713 segments from TD children, respectively.

### 2.6 EEG connectivity matrices

To transform the pre-processed EEG segments into graph-like matrices, we employed three different connectivity measures in this study: the Pearson correlation coefficients (PCC), phase locking value (PLV), and Rho index (RHO). These FC measures, expressed in terms of signal correlation and phase synchronization, are further detailed as follows.

(1) The PCC measures the linear correlation between two signals (Cohen et al., 2009). It ranges between -1 and 1. The value of -1 means a perfect negative linear correlation between the two signals, 0 means the two EEG signals are not linearly correlated, and 1 represents the perfect positive linear correlation between the signals. Let 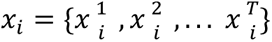denote an EEG signal of the *i*-th electrode, where *T*is the time length of the signal. The PCC of the two signals 𝑥_𝑖_ and 𝑥_𝑘_ is calculated as:

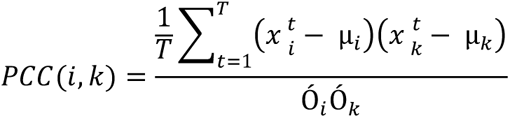

Where *μ* and *Ó* are the mean and standard deviation of the signal, respectively.

(2) The PLV is a prevalent neuroimaging metric for quantifying phase synchronization between two signals. It estimates how the relative phase is distributed over the circle (Lachaux et al., 1999). PLV values range from 0 to 1, with higher values indicating more robust phase synchrony between the two signals at the specified frequency band. A PLV of 0 indicates no phase locking, while a PLV of 1 indicates perfect phase locking. It can be quantified using the following formula:

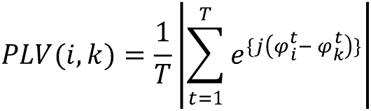

Where 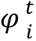is the phase of the signal at time *t*.

(3) The RHO index is measured based on the Shannon entropy (Tass et al., 1998). It quantifies the deviation of the distribution of the cyclic relative form from the uniform distribution. The RHO index ranges from 0 to 1, where a value of 0 indicates no synchronization, while a value of 1 indicates perfect synchronization. It can be calculated as follows:

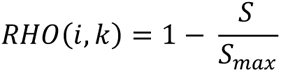

Where 𝑆_𝑚𝑎𝑥_ is the maximal entropy (that of uniform distribution), i.e., the logarithm of the number of bins in the histogram, and *S*is the entropy of the distribution of Δ𝜑_𝑟𝑒𝑙_ (𝑡):

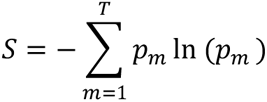

where, 𝑝_𝑚_ is the probability of finding Δ𝜑_𝑟𝑒𝑙_(t) in the *m*-th bin.

In our study, we implemented these FC measures for each segment and frequency band in all experimental conditions. The computation of FC principally utilized the HERMES brain connectivity toolbox (https://hermes.med.ucm.es), operating on the MATLAB R2018b environment (MathWorks Inc., Natick, MA, USA). Connectivity features were ascertained for each pair of EEG electrodes, culminating in an extensive feature set representable as a connectivity matrix. Within this matrix, the value at the (*i*, *j*) position corresponds to the connectivity measure between the EEG signals from the *i*-th and *j*-th electrodes (see Figure 3). This matrix serves analogously to an adjacency matrix in graph theory, where the nodes correspond to the EEG electrodes, and the connectivity values give the edge weights.

**Figure 3.**
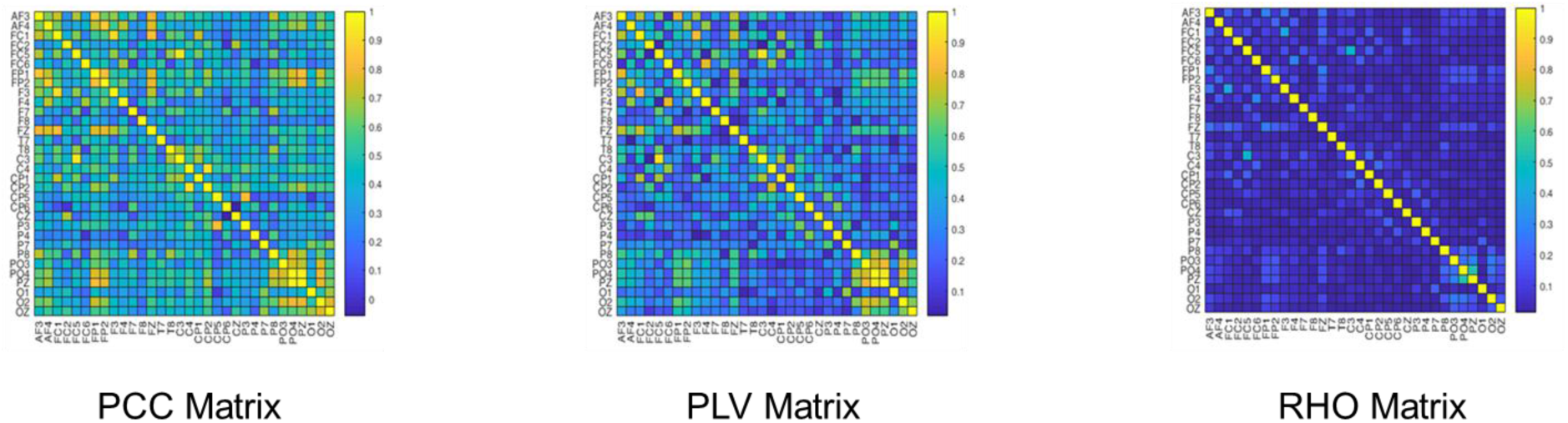
FC matrices for a segment in the beta band from a child with developmental dyslexia. Notes: Each matrix visualizes the connectivity between pairs of EEG electrodes. The intensity of color in each cell represents the strength of connectivity between the corresponding pair of electrodes.

### 2.7 CNNs classifier

In our study, we utilized the CNNs model that comprised multiple layers: two convolutional layers with ReLU activation functions, two max-pooling layers, two batch normalization layers, two dropout layers, a flatten layer a fully connected layer, and an output layer (as detailed in Table 2). The first and second convolution layers utilized a 3 × 3 convolution kernel with 32 filters, followed by a MaxPooling2D layer. This layer served to compress spatial dimensions while retaining vital information. Each max-pooling layer was succeeded by a BatchNormalization layer to normalize the activations, simplifying the learning process and enhancing the speed and performance of the model (Jung et al., 2019). A Dropout layer was applied following the second batch normalization layer, randomly deactivating a fraction of input units during training to help prevent over-fitting (Wu & Gu, 2015). The 2D arrays were converted into a 1D array by a Flatten layer before being fed into the Dense layer, a fully connected layer with 128 neurons. To further guard against over-fitting, we applied another Dropout layer after this Dense layer. Finally, the output layer, another Dense layer equipped with a single neuron, was used for the binary classification task.

We compiled our CNNs model using the Adam optimizer, which stands for Adaptive Moment Estimation (Mehta et al., 2019). This method calculates adaptive learning rates for each parameter, making it a favored choice for deep learning applications. We utilized binary cross-entropy as the loss function, which is apt for binary classification tasks. This function measures the difference between predicted and actual labels. The model’s performance was assessed using the accuracy metric, which compares the model’s predictions with the actual values and calculates the percentage of correct predictions. The model was designed to iterate over the data for 100 epochs.

The CNNs model and classification process were implemented in Python using TensorFlow. All the training, validation, and testing processes were carried out on an Intel(R) Xeon® Bronze 3204 CPU with six cores, supported by an NVIDIA Quadro P2000 GPU.

**Table 2.** Convolition Neural Networks architecture.

| Layer | Type | Output shape | Kernel size |
| --- | --- | --- | --- |
| 1 | Convolution | $32 \times 32 \times 32$ | $3 \times 3$ |
| 2 | Max-pooling | $16 \times 16 \times 32$ | $2 \times 2$ |
| 3 | Batch_normalization | $16 \times 16 \times 32$ | - |
| 4 | Convolution | $16 \times 16 \times 64$ | $3 \times 3$ |
| 5 | Max-pooling | $8 \times 8 \times 64$ | $2 \times 2$ |
| 6 | Batch_normalization | $8 \times 8 \times 64$ | - |
| 7 | Dropout | $8 \times 8 \times 64$ | - |
| 8 | Flatten | 4096 | - |
| 9 | Dense | 128 | - |
| 10 | Dropout | 128 | - |
| 11 | Dense | 1 | - |

### 2.8 CNNs training and cross-sample validation

Before initiating the training process, we divided the FC matrices of each experimental condition into two independent samples based on the participant’s ID. We allocated 80% of the data to Sample 1 and 20% to Sample 2, ensuring that both samples maintained the same ratio of dyslexia segments to control segments.

Sample 1 was used to train and validate the CNNs model. We employed a 5-fold cross-validation method to evaluate the model’s performance. This involved splitting Sample 1 randomly into five equal parts or folds. The model was trained on four folds and validated on the remaining one. This procedure was repeated five times, with each fold used as the validation set exactly once. The final accuracy score was computed as the average of the five validation scores.

After training and validating the CNNs model, we saved the best model for the five-fold cross-sample validation. We then of applied the best-performing frequency band’s model to Sample 2 to test its effectiveness in the cross-sample validation. Moreover, to confirm the robustness of our results, a permutation test was conducted 1,000 times to establish if the cross-sample validation accuracy was significantly higher than the chance level.

### 2.9 Identifying the discriminative FCs contributing to the CNNs classification

Drawing inspiration from the hierarchical organization of features in CNNs (Drozdzal et al., 2016; Nguyen et al., 2017), we posited that the features that help distinguish children with DD from TD peers might also represent the different brain patterns between these two groups. Therefore, we directly compared children with DD and their TD peers in Sample 1 using the best-performing FC indices of all frequency bands. We conducted independent-sample *t*-tests on each FC index to identify the connectivity values that were significantly higher in the DD group compared to the TD group (DD_TD) and those that were significantly higher in the TD group compared to the DD group (TD_DD). The FDR correction method at a *q* < .01 level was used to mitigate the issue of multiple comparisons. For a more precise visualization of the discriminative FCs, we used the BrainNet viewer (https://www.nitrc.org/projects/bnv/)on MATLAB (MathWorks Inc., Natick, MA, USA).

### 2.10 Exploring the relationship between identified discriminative FC and Chinese word reading performance

Further, we aimed to explore the relationship between discriminative FCs and Chinese word reading performance within the DD and TD groups, respectively. To accomplish this, we calculated each participant’s average discriminative FC values. Subsequently, we performed permutation-based correlation analyses (number of permutations = 10,000) to assess the correlations between these mean FC values and the measures of Chinese word reading accuracy and fluency within the DD and TD groups, respectively.

## 3. Results

### 3.1 Convolutional neural networks performance

We used the extracted FC indices of different frequency bands and CNNs to classify children with DD and TD peers in all four experimental conditions. Table 3illustrates the classification accuracy in these conditions for each frequency band. The results indicate that the beta-based FC indices consistently surpassed other indices in identifying DD among all frequency bands, with a mean classification accuracy of 90.96% across the four experimental conditions. Conversely, the delta-based FC indices had the lowest effectiveness in distinguishing DD, achieving a mean accuracy of 67.88% across all experimental conditions. Interestingly, eyes-open and eyes-closed FC indices (with mean accuracies of 79.55% and 79.52%, respectively) were more successful in identifying DD compared to the verbal one-back and verbal two-back conditions (with mean accuracies of 70.07% and 70.45%, respectively). Among different types of FC measures, the RHO index was particularly notable as it was the most effective in identifying DD, showing superior performance in most experimental conditions and frequency bands. Specifically, the beta band RHO index in the eyes-open condition achieved the highest classification accuracy, reaching 97.66% (±0.31%), among all FC indices. Figure 4 provides detailed learning curves for the eyes-open beta band RHO index.

**Figure 4.**
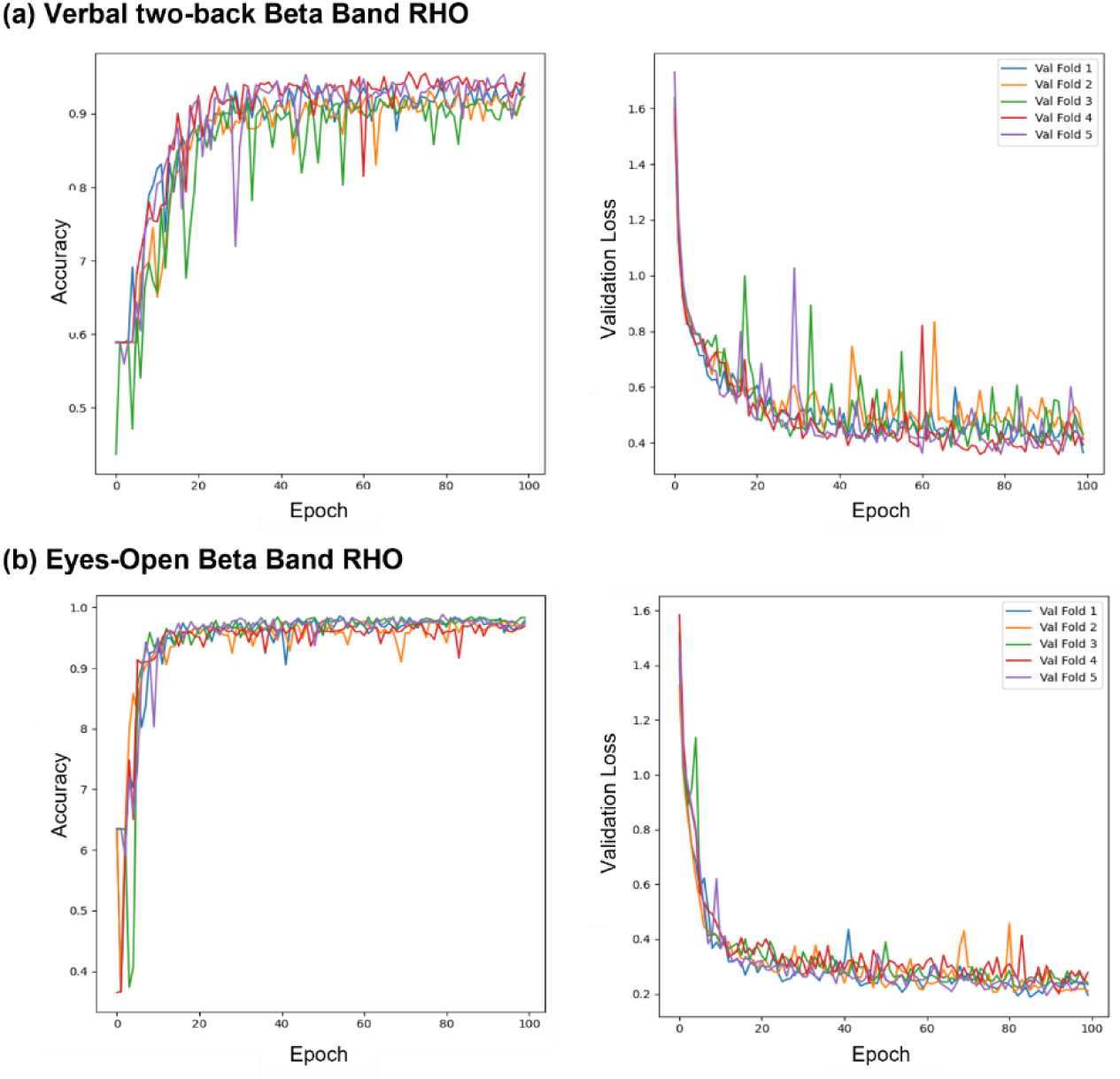
Learning curves of the CNN model trained with RHO connectivity. Notes: The learning curves of the CNN model were trained using 5-fold cross-validation with RHO connectivity features within the beta band. (a) Shows the learning curves for the eyes-open resting-state condition, and (b) shows the learning curves for the verbal two-back task condition. Validation accuracy and validation loss are plotted across epochs to assess model performance and generalization.

**Table 3.** CNNs five-fold classification accuracy.

| Condition | FC | Delta | Theta | Alpha | Beta |
| --- | --- | --- | --- | --- | --- |
| Eyes-open | PCC | 66.01±1.31 | 73.62±1.19 | 75.08±1.27 | 91.39±1.49 |
|  | PLV | 70.24±0.73 | 74.63±1.37 | 75.10±0.55 | 95.19±0.68 |
|  | RHO | 75.08±1.13 | 80.23±2.13 | 80.41±1.28 | 97.66±0.31 |
| Eyes-closed | PCC | 67.06±0.70 | 71.87±1.22 | 76.98±1.81 | 91.24±0.38 |
|  | PLV | 69.91±1.29 | 75.33±0.51 | 77.01±0.85 | 94.76±0.91 |
|  | RHO | 72.07±2.15 | 79.68±1.27 | 81.26±1.44 | 97.07±0.62 |
| Verbal one-back | PCC | 72.65±2.24 | 56.46±2.53 | 65.35±1.32 | 81.69±0.40 |
|  | PLV | 61.94±0.73 | 65.15±0.99 | 64.45±1.53 | 88.07±0.62 |
|  | RHO | 61.01±1.67 | 64.65±3.03 | 67.83±2.92 | 91.57±1.46 |
| Verbal two-back | PCC | 74.93±1.81 | 57.25±2.27 | 63.27±1.72 | 81.86±1.39 |
|  | PLV | 60.77±0.91 | 65.44±1.33 | 66.38±1.02 | 88.70±2.23 |
|  | RHO | 62.88±1.31 | 66.53±0.81 | 65.08±0.87 | 92.29±1.18 |

**Notes:** Classifying typically developing (TD) and developmental dyslexia (DD) groups with various FC indices of different frequency bands in four experimental conditions.

### 3.2 Cross-sample validation performance

We further conducted cross-sample validation on the best-performing beta band indices to test whether the trained CNNs models with high classification accuracies could be generalized to screen DD in the independent Sample 2. The validation accuracies remained above chance level across all experimental conditions. As shown in Table 4, the beta-band RHO index in the eyes-open condition achieved the highest cross-sample validation accuracy of 69.95% among the various FC indices.

**Table 4.** Cross-sample validation accuracy.

| Condition | FC | Cross-sample validation accuracy |
| --- | --- | --- |
| Eyes-open | PCC | 68.21** |
|  | PLV | 63.68** |
|  | RHO | 69.95** |
| Eyes-closed | PCC | 60.61** |
|  | PLV | 64.44** |
|  | RHO | 65.96** |
| Verbal one-back | PCC | 58.95** |
|  | PLV | 61.15** |
|  | RHO | 59.31** |
| Verbal two-back | PCC | 56.60** |
|  | PLV | 61.76** |
|  | RHO | 60.38** |

**Notes:** Cross-sample validation accuracy for predicting typically developing (TD) and developmental dyslexia (DD) groups using the pre-trained CNN model with beta-band functional connectivity (FC) features across different experimental conditions. \*\* Significant higher than the chance level ( $p < .001$ ).

### 3.3 Identifying discriminative FCs

The CNNs classification results revealed that the beta band RHO index in the eyes-open condition was particularly effective in distinguishing children with DD from TD peers, achieving an impressive accuracy of 97.66%. To further probe into the discriminative FCs, we conducted independent-sample *t*-tests on the eyes-open beta band RHO index, with FDR correction at *q*< .01 level, within Sample 1. Figure 5(d) shows that a significant number of discriminative FCs existed between the DD and TD groups, including scenarios where TD children’s FC values were stronger than those of children with DD and vice versa. More specifically, TD children exhibited stronger FCs between the temporal and parietal scalp areas, the temporal and frontal-central scalp areas, and the central and central-parietal scalp areas compared to children with DD. In contrast, children with DD exhibited stronger discriminative FCs in the frontal-central and frontal scalp areas, the frontal-parietal and anterior-frontal scalp areas, and the frontal-central and parietal scalp areas compared to TD children.

In addition to the beta band RHO index, our study also tried to identify discriminative FCs of the other frequency bands in the eyes-open condition. As shown in Figure 5, discriminative FCs where TD children exhibited stronger FCs than children with DD were primarily located in the central and central-parietal scalp areas and the central and occipital scalp areas for the other three frequency bands. Conversely, for these frequency bands, children with DD displayed stronger FCs than TD children in the frontal and frontal-central scalp areas and the frontal and anterior-frontal scalp regions. These frequency band patterns partially align with the discriminative FC pattern observed in the beta band. When comparing different frequency bands, we observed that the FC pattern of the beta band exhibited more robust discrimination between the DD and TD groups than other frequency bands, no matter whether the DD group displayed stronger FCs than the TD group or vice versa.

**Figure 5.**
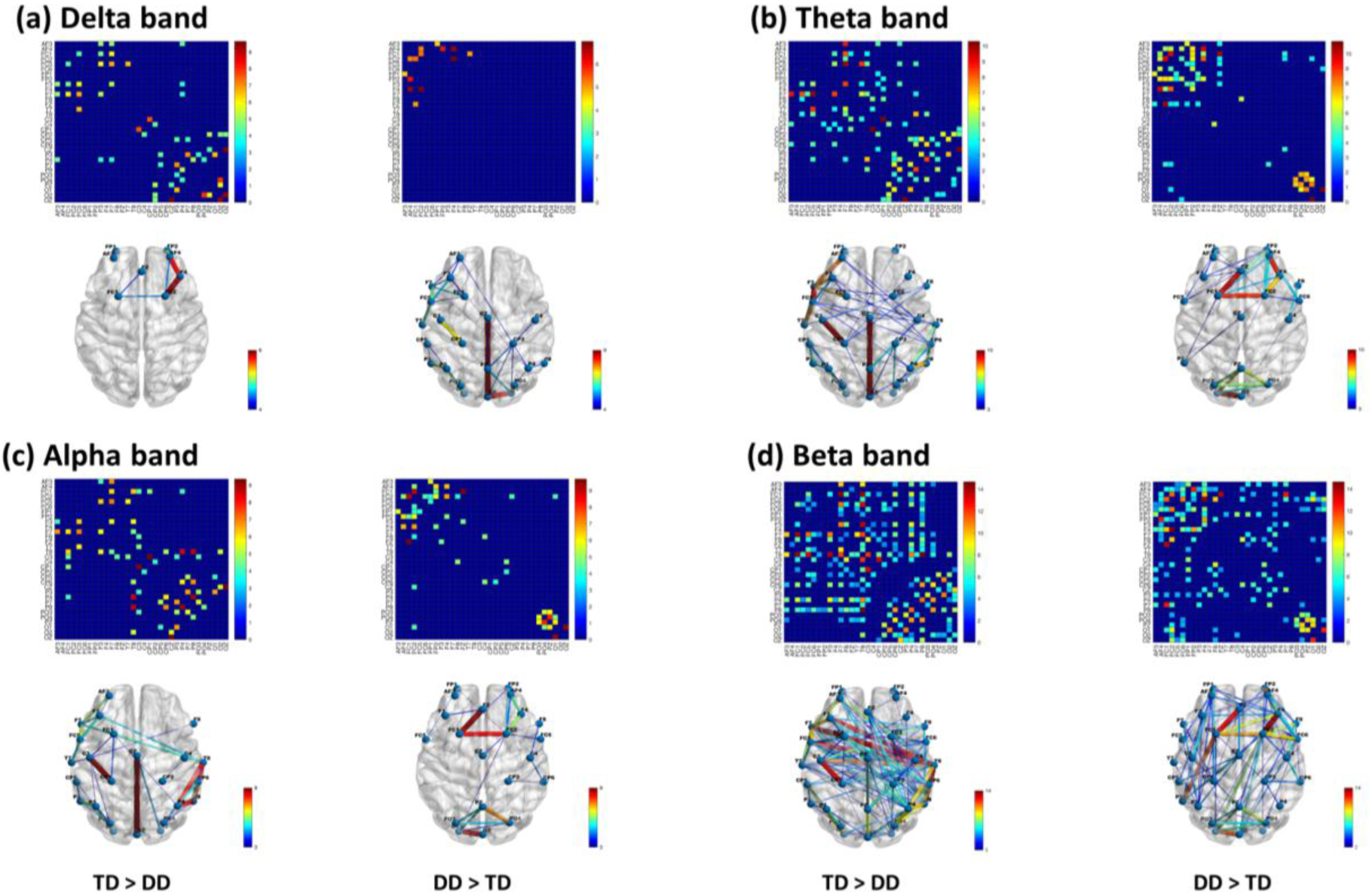
Discriminative FCs between typically developing (TD) children and children with developmental dyslexia (DD) in the eyes-open condition. Notes: The color bar represents the *t*-value from each comparison of RHO indices between the TD and DD groups within Sample 1, with FDR correction at *q*< .01 level. Abbreviations: DD (Developmental Dyslexia), TD (Typically Developing).

### 3.4 Linking discriminative FCs to Chinese word reading performance

Our study also aimed to explore the association between discriminative FCs and Chinese word reading abilities. To this end, we conducted separate correlation analyses within the DD and TD groups, assessing the association between the average discriminative FCs derived from the eyes-open beta band RHO index of each participant and their respective Chinese word reading abilities. Interestingly, significant correlations were observed within the DD group only. In this group, where discriminative FCs were notably stronger compared to the TD group, the average discriminative FCs were significantly negatively correlated with both Chinese word reading accuracy (*r*

= -.26, *p*< .05) and fluency (*r*= -.28, *p*< .05). These findings suggest that within the DD group, higher FC values are associated with lower performance in both reading accuracy and fluency.

## 4. Discussion

The current study integrated EEG-based FC measures both at resting- and task-states with CNNs for effective screening and identification of behaviorally relevant neural biomarkers in Chinese DD. Our CNNs classification results showed that beta band FC indices consistently outperformed those of other frequency bands. These results suggest a high potential for beta band FC indices in screening for DD. We further analyzed the generalizability of our model through cross-sample validation. Although the cross-sample accuracy was lower than the within-participant classification accuracy, it consistently exceeded the chance level, demonstrating the generalizability of our trained CNN model for DD screening. Furthermore, we identified distinctive FCs contributing to DD classification, offering potential neural biomarkers for Chinese DD. Lastly, we probed the correlation between the identified discriminative FCs and reading abilities, forging a connection between our neuroimaging findings and behavioral outcomes.

Our CNNs classification results consistently demonstrated that FC indices in the beta band performed superior to delta, theta, and alpha bands across all experimental conditions. This aligns with existing research that has identified atypical beta band activity in children with DD in comparison to TD children (Cainelli et al., 2023; Penolazzi et al., 2010; Spironelli et al., 2008). Additionally, an EEG connectivity study also found increased beta band coherence in the frontal scalp area when comparing DD children to TD peers (Arns et al., 2007). Similarly, EEG graph studies employing FC measures have unveiled abnormal beta band network patterns in DD children when contrasted with TD children (Taskov & Dushanova, 2021; Xue et al., 2020). These findings suggested that the abnormalities in the beta band could differentiate the neural patterns between children with DD and TD children, thus contributing to higher classification accuracy. Previous studies also found that the beta band activities were related to executive functions (Basharpoor et al., 2021; Wang et al., 2022), as well as visuomotor integration (Kilavik et al., 2013; Lin et al., 2012). The link between Chinese word reading and executive functions and visuomotor integration was reported in many studies (Chung & McBride-Chang, 2011; McBride-Chang et al., 2011; Liu, Chung, & Fung, 2019). Taken together, these studies suggested that the distinctiveness of beta band FC indices in our CNNs classification results might have arisen from the involvement of anomalous executive functions and visuomotor integration processes among Chinese children with DD. This evidence collectively supports the effectiveness of beta band FC indices in distinguishing children with DD from their TD peers, emphasizing the potential of beta band activities as a neural biomarker for Chinese DD.

The CNNs classification results further suggested the superior efficacy of resting-state FC indices in the beta band over *n*-back task-state FC indices in screening children with DD. Prior research indicated that resting-state brain activity can provide a broader perspective of neural interactions and illuminate the fundamental organization of the brain’s functional system (Deco et al., 2011; Fox & Raichle, 2007). This more comprehensive view of neural interaction could be particularly beneficial in identifying unique brain patterns that differentiate children with DD from TD children,. In contrast, the verbal *n*-back task primarily evaluates the neural processes associated with managing working memory demands (Deco et al., 2011; Fox & Raichle, 2007), which may correlate with a more specific aspect of reading abilities. Hence, this task might not encompass all the crucial information necessary for differentiating between DD and TD children. Nonetheless, it is conceivable that task-state EEG data from alternative reading-related tasks could yield better predictions for dyslexia. Furthermore, our results indicate that resting-state EEG outperforms the *n*-back task-state EEG, suggesting that DD may stem from mechanisms involving more domain-general rather than domain-specific processing. Given that DD is associated with deficits in phonological (Ramus, 2003), orthographic (Su et al., 2015), morphological (Breadmore & Carroll, 2016), and visual-auditory processing (Schulte-Körne & Bruder, 2010), focusing solely on working memory may not suffice for accurate differentiation. These findings support the notion that DD is a multifaceted disorder characterized by domain-general deficits, underscoring the importance of considering a broad spectrum of cognitive and neural processes in its assessment and diagnosis.

Among the different FC measures, the RHO indices outperformed PCC and PLV in screening children with DD most of the time. While PCC primarily captures the linear correlation between two EEG signals (Guevara & Corsi-Cabrera, 1996), the PLV assesses the consistency of the phase relationship over time (Lachaux et al., 1999). The RHO index, however, takes a more nuanced approach by quantifying the variability in phase synchronization. It evaluates the degree of deviation from a uniform distribution, which can reflect more complex patterns of neural connectivity (Tass et al., 1998). Because phase synchronization measures like PLV and RHO indices can detect intricate neurophysiological information beyond simple linear correlations, they are potentially more effective for identifying DD. The RHO index, in particular, demonstrated a slightly higher accuracy in distinguishing between the DD and TD groups compared to the PLV. This increased accuracy from RHO might have stemmed from its ability to account for the entropy within phase relationships, giving it a heightened sensitivity to complex and subtle phase synchronization patterns. Such sensitivity may be crucial for identifying neurophysiological differences between the DD and TD groups that the PLV might miss, as the latter primarily gauges the consistent phase locking between signals. Furthermore, Wang et al. (2020)have shown that the RHO index achieved the highest accuracy for EEG biometric identification compared to other connectivity measures evaluated, suggesting that the RHO index may have proficiency in EEG-based FC classification tasks. In addition, the enhanced performance of the RHO index might also be attributed to the CNNs architecture used in our study. This advanced analytical approach is likely more capable of discerning the subtle differences in EEG signals between children with DD and their TD peers, particularly when utilizing the RHO index.

In contrast to previous studies that primarily focused on classifying children with DD at the within-participant level, the current study extended the performance evaluation of our trained CNNs to an independent sample. Poldrack et al. (2020)highlighted the need for a strict separation between training and test data to avert information ’leakage’ that could artificially boost accuracy estimates. Furthermore, exclusive reliance on k-fold cross-validation within the original dataset may not ensure data independence, particularly if training and validation datasets encompass overlapping data from the same participants. Consequently, cross-sample validation becomes essential to verify our model’s predictive accuracy and to mitigate over-fitting risks. Our cross-sample validation results showed that the classification accuracy of all FC indices in the beta band surpassed the chance level. However, there was a marked decrease in cross-sample classification accuracy compared to the within-participant classification accuracy. The beta band RHO index under the eyes-open condition yielded a cross-sample validation accuracy of approximately 70%. This performance lagged considerably behind the within-participant classification, which boasted a classification accuracy of about 98%. This discrepancy indicates that our trained CNNs classifier could not generalize effectively across participants for the classification of DD. This could be attributed to the considerable variability both within and between participants. DD is a multifaceted learning disorder presumably with a neurological basis (Beelen et al., 2019; Yan et al., 2021). Even though children were identified as having DD, the multitude of potential cognitive and neurological causes could lead to substantial variability between participants, thereby decreasing the generalizability of our trained CNNs. Additionally, the significant decline in cross-sample accuracy raises the possibility of overfitting, indicating that our CNNs model might have been overly optimized for the training data to the detriment of its performance on independent datasets. This issue may also hint at an insufficient separation between training and testing data within the five-fold cross-validation, suggesting the chance of information leakage (Poldrack et al., 2020). Addressing these challenges will be crucial in refining the application of CNNs for the screening of DD.

Further, we identified the discriminative FCs that significantly contributed to screening children with DD. We observed more discriminative beta band FCs than other frequency bands in scenarios where FCs in the TD group were stronger than those in DD and vice versa. Moreover, the beta band FCs exhibited more pronounced peak *t*-values in these scenarios than FCs in the other three frequency bands. These findings hint that these discriminative FCs could explain the highest classification accuracy in the beta band in the CNNs model due to more and stronger discriminative FCs. In addition, we found the most discriminative FCs that the TD group is stronger than the DD group were primarily found between the temporal and parietal scalp areas, the temporal and frontal-central scalp areas, and the central and central-parietal scalp areas. Our findings partially align with previous research, which reported that during task-state EEG, TD children demonstrated stronger beta band connections, particularly between the T5-T6 and Cz-C3 electrodes, compared to children with DD (Seshadri et al., 2022). Moreover, the temporal and parietal regions have been suggested to be associated with the reading process (Cohen et al., 2000; Simos et al., 2002). Therefore, the increased connectivity in the temporal, parietal, and central areas among TD children compared to DD children may correlate with the reading process dysfunctions in children with DD. In contrast, the most discriminative FCs that the DD group is stronger than the TD group were primarily found in the frontal-central and frontal scalp areas, the frontal-parietal and anterior-frontal regions scalp areas, and the frontal-central and parietal scalp areas. These findings align with previous studies showing increased connectivity in children with DD and adults with dyslexia during reading and tasks between multiple frontal and parietal areas (Finn et al., 2014; Hosseini et al., 2013, Richards & Berninger, 2008; Schurz et al., 2015). Besides, Žarić et al. (2017) discovered increased connectivity between the right and left frontal sites during visual word false font processing. Furthermore, previous studies suggested that increased frontal activation could represent a compensatory mechanism, particularly in adults, associated with articulatory processes during visual word processing (Shaywitz & Shaywitz, 2005; Richlan, Kronbichler, & Wimmer, 2011). Therefore, more robust connectivity between frontal scalp areas in children with DD may reflect more effortful strategies and compensatory mechanisms engaged during visual word processing.

Lastly, our study separately examined the relationship between discriminative FCs and Chinese word reading performance in children within the DD and TD groups, respectively. Our analyses revealed that within the DD group, there was a significant negative correlation between discriminative FCs—stronger in this group compared to the TD group—and both reading accuracy and fluency. Specifically, these correlations were quantified as *r*= -.26, *p*< .05 for reading accuracy, and *r*= -.28, *p*< .05 for fluency, suggesting that children with DD exhibit particular neural connectivity patterns that adversely affect their reading abilities. These findings underscore the complex and nuanced relationship between neural connectivity and reading performance in the DD group. However, it is important to note that the correlation coefficients, while significant, were relatively modest. This indicates that while discriminative FCs capture some critical aspects of the neural underpinnings of reading, they do not encompass all the factors that contribute to reading performance. This highlights the possibility that multiple neural pathways or additional external factors may influence reading abilities, suggesting avenues for further research.

Despite our CNNs model demonstrating remarkable accuracy in screening children with DD within individual participants, several methodological limitations must be determined. Firstly, a notable decrease in classification accuracy was observed when the model was applied to independent samples, in which data segments span multiple participant levels. This drop in accuracy may be ascribed to substantial variability among participants. Given the diminished cross-sample validation accuracy, which may fall below the threshold of clinical acceptability, the current model faces challenges for practical deployment in clinical settings. Future research could address this issue by training the CNN classifier on larger sample sizes, thus incorporating more participants. This approach would likely diminish the impact of individual variability and enhance the model’s generalizability. Secondly, our study utilized FC measures at the scalp level, allowing us to interpret dysfunctions as they manifest at the EEG electrode level. However, this approach limits our ability to comprehensively understand brain connectivity abnormalities at the brain area level. Future EEG studies aiming to screen children with DD and identify potential biomarkers should consider adopting FC measures based on EEG source localization (Lei et al., 2022; Zhang et al., 2021). This approach could improve the interpretation of identified brain biomarkers and enable cross-modularity validation with other types of neuroimaging data, such as fMRI. Lastly, while this study utilized EEG data from verbal working memory tasks for screening children with DD, the classification performance of task-state EEG data was inferior to resting-state EEG. The verbal *n*-back tasks might have captured a limited amount of group difference only. Future studies might explore incorporating other types of reading-related tasks and concurrent EEG recording to enhance the screening process for children with DD.

Overall, the current study employed EEG data in conjunction with deep learning techniques to screen Hong Kong Chinese children with DD, achieving an outstanding classification accuracy of approximately 98% at the within-participant level and a respectable accuracy of about 70% at the cross-participant level. Furthermore, we identified potential biomarkers - the discriminative FCs - that could facilitate the screening of DD and explored their relationship with reading ability. The findings of this study could contribute to the development of a neuroimaging-based screening tool for DD in Chinese children and may also be adaptable to other languages and writing systems. Therefore, this research has scientific implications for understanding the neural underpinnings of DD and holds potential practical applications in early interventions and educational strategies for children with DD.

## Acknowledgements

This research was supported by the Health and Medical Research Fund (HMRF) of the Food and Health Bureau, Hong Kong (04152496), a Direct Grant from the Faculty of Social Sciences at the Chinese University of Hong Kong (4052177), and the General Research Fund (GRF) of the Research Grants Council of Hong Kong (RGC-GRF 14600919).

## Declaration of Generative AI and AI-Assisted Technologies in the Writing Process

During the preparation of this work, the authors used OpenAI’s ChatGPT to assist with language refinement and editing for clarity and flow. After using this tool, the authors reviewed and edited the content as needed and take full responsibility for the content of the published article.

